# NBDC Data Tools: Building Analysis-ready Datasets From ABCD and HBCD Releases in R and Python

**DOI:** 10.64898/2026.11.02.763757

**Authors:** Le Zhang, Biplabendu Das, Olivier Celhay, Calen J. Smith, Laura R. Ziemer, Anders M. Dale, Janosch Linkersdörfer

## Abstract

The NIH Brain Development Cohorts (NBDC) Data Hub provides access to large-scale longitudinal datasets, including the Adolescent Brain Cognitive Development (ABCD) Study® and the HEALthy Brain and Child Development (HBCD) Study. The studies release their tabulated data as hundreds of tables with tens of thousands of variables. Turning these files into a dataset for a particular analysis poses substantial informatics challenges, especially when data preparation and analysis workflows must be reproducible and reusable for new releases. We therefore developed *NBDC Data Tools*, a suite of open-source packages for preparing NBDC data in local or other approved computing environments. *NBDCtools* is the core R package for dataset construction and processing. The Python package *nbdctools* either calls the R workflow or assembles datasets natively in Python, and the R package *NBDCtoolsData* provides the release metadata support for both *NBDCtools* and *nbdctools*. Users define a dataset in code by naming the study, the release, and the variables or tables they need. The software then uses the standardized metadata released by the studies to locate the relevant files, join tables on the participant and session identifiers, convert data types, attach labels, and optionally recode non-response codes as missing values. Because the dataset definition is a short script that runs on the original release files, it can be rerun, shared with collaborators, and reported in publications. *NBDC Data Tools* complements the web-based data platforms of the NBDC Data Hub by providing a reproducible, scriptable path from approved data access to analysis-ready data for statistical analysis.

## Introduction

Large-scale longitudinal cohort studies have become foundational resources for understanding human brain development, behavior, and health across the lifespan [1–3]. The NIH Brain Development Cohorts (NBDC) Data Hub (https://www.nbdc-datahub.org/) represents a significant advancement in open science, providing researchers worldwide with access to comprehensive, curated datasets from large longitudinal studies, including the Adolescent Brain Cognitive Development Study (ABCD Study®) and the HEALthy Brain and Child Development (HBCD) Study [4]. The ABCD Study® is the largest longitudinal study of brain development and child health in the United States [5,6]. The HBCD Study, which follows families from pregnancy through early childhood, is the largest longitudinal study of early brain and child development in the United States [3].

Despite their scientific value, preparing NBDC data for analysis in a declarative and reproducible manner constitutes substantial informatics challenges [4,6]. Tabulated data from NBDC datasets are curated and released in a high-dimensional hierarchical structure, where thousands of variables are partitioned across disparate data tables corresponding to specific measurement domains and instruments. In practice, researchers frequently need to analyze different measures across multiple domains, such as neuroimaging, neurocognition, mental health, or substance use, with sociodemographic and other participant information. Constructing custom analysis-specific datasets containing all the variables needed for a given analysis requires complex relational joins, systematic handling of missingness, and applying explicit data type conversions. These steps require extensive knowledge of data curation standards and the structure of each release, as well as advanced data-processing skills [7,8].

In many projects, these challenges are addressed either through repeated creation and download of custom datasets through web-based interfaces and/or through user-specific *ad hoc* preprocessing scripts [9]. Within the NBDC Data Hub, the two main access points are the Data Exploration & Analysis Portal (DEAP, https://nbdc.deapscience.com/) and the NBDC Data Access Platform (DAP, https://nbdc-datashare.lassoinformatics.com/), which support exploring, querying, and downloading data and creating custom datasets through web-based workflows [4]. These web platforms provide important infrastructure for accessing and exploring data from NBDC studies and a convenient way to download data for analysis, but are less suited for iterative, fully reproducible workflows in which datasets must be rebuilt repeatedly to accommodate revised variable sets, new covariates, updated study releases, or alternative inclusion criteria [9]. Local *ad hoc* scripts avoid the manual steps but are often difficult to standardize, audit, and reproduce across analysts and projects [10]. A script shared with a collaborator also tends to fail because it starts from an intermediate data file that only its author has.

Software developed for other cohort resources has shown how much resource-specific tooling can mitigate complex preprocessing burdens. For the UK Biobank, for example, the R package *ukbtools* and the Python toolkit *FUNPACK* facilitate dataset assembly by parsing UK Biobank releases alongside field-level metadata, but they are built around Biobank’s own file structures and metadata standards [11,12]. Despite these advances, preparing NBDC cohort data lacks a programmatic framework that leverages standardized metadata to construct analysis-ready datasets from explicit code definitions, allowing researchers to re-execute workflows on original tabular releases as analytic requirements or dataset versions evolve.

To resolve these challenges, we developed *NBDC Data Tools*, a suite of open-source software that enables automated, metadata-guided assembly of NBDC datasets directly from tabulated release files. Centered on the R package, *NBDCtools* reads the per-table files of a downloaded release, selects and joins the requested variables and tables, and applies processing steps such as factor conversion and labeling. *NBDCtoolsData* supplies the metadata for each supported release, and the Python package *nbdctools* gives Python users access to the same workflow. Dataset assembly and preprocessing run in one scripted call, so the code that defines a dataset also documents how it was derived. In this paper, we describe the design and implementation of the suite, illustrate its use with a cross-domain example in R and Python, show how a dataset definition can be reused, and report benchmarks of runtime and memory use.

## Methods & Implementation

### Software suite overview

*NBDC Data Tools* is designed as a suite of software for reproducible preparation of NBDC tabulated data (Fig. 1A). The workflow requires locally downloaded tabulated release data files in release format (one file per table) and associated metadata, including:

a. data dictionaries (*dd*, tables that define variable description, type, and other information);
b. levels tables (*lvl*, tables that map numeric categorical codes to their human-readable category labels);
c. sessions (*ses*, metadata that tracks event names, visit timing, and wave structures);
d. shadow matrices (HBCD only; tables that have the same dimensions as the main data table and encode specific missingness reason codes for every cell value).

**Figure 1.**
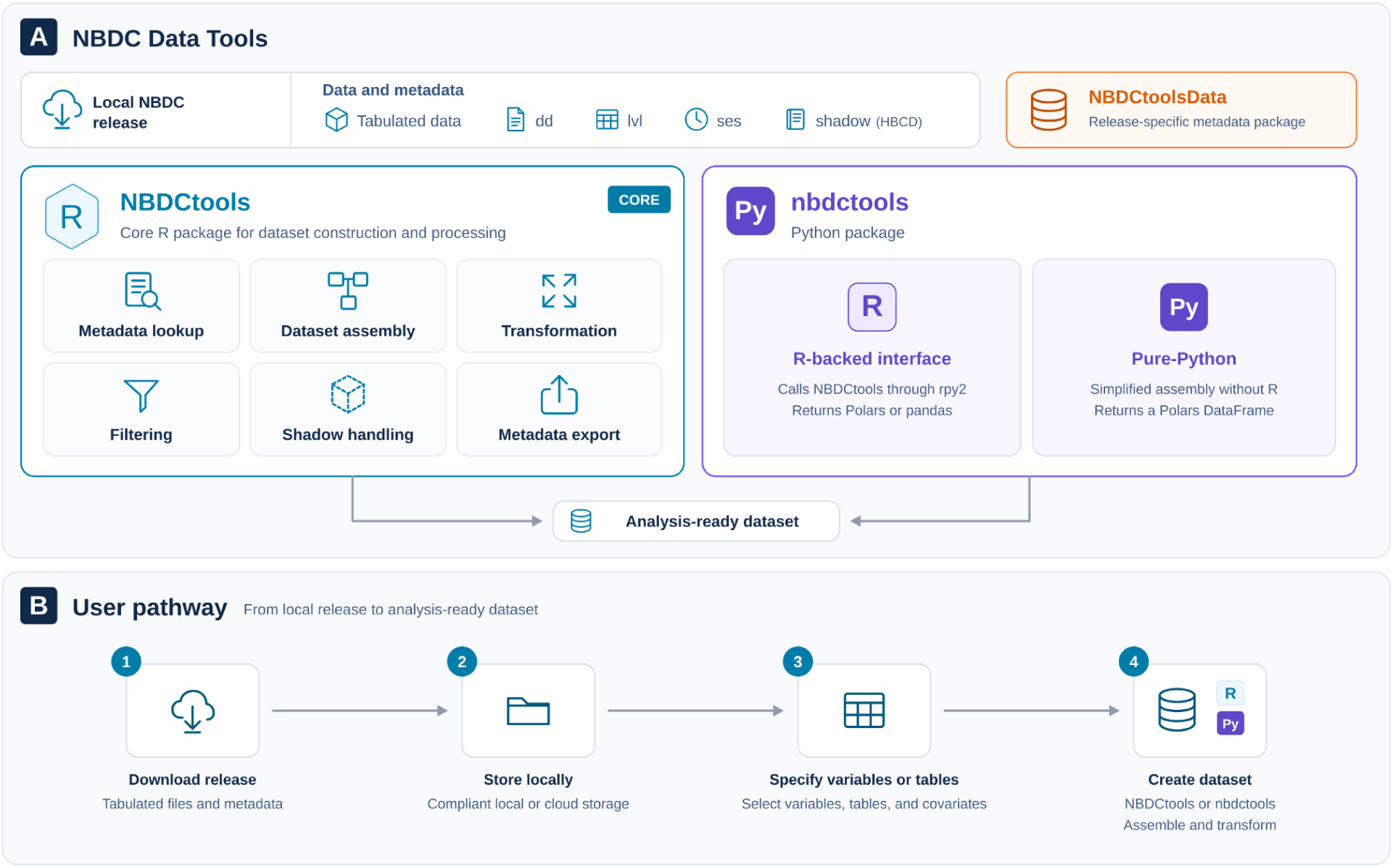
Overview of *NBDC Data Tools* and the user workflow. (A) The suite connects locally downloaded NBDC tabulated data and metadata with the core R package *NBDCtools*, the Python package *nbdctools*, and the metadata package *NBDCtoolsData*, which together produce analysis-ready datasets. (B) Users download a tabulated release, store the files locally, specify the variables or tables they need, and create the dataset with *NBDCtools* or *nbdctools*.

The core R package NBDCtools uses these inputs for metadata lookup, dataset assembly, transformation, filtering, shadow matrix handling, and metadata export.

Two companion packages complete the suite. *NBDCtoolsData* provides the release-specific metadata as an R data package. It is updated with each data release, so *NBDCtools* itself does not need to change when a new release appears. The Python package *nbdctools* can either call the R implementation of the dataset-construction workflow or assemble datasets with a simplified pure-Python implementation, which helps users who cannot or prefer not to install R (Fig. 1A, Table 1).

**Table 1.** Packages in *NBDC Data Tools* and their roles

| **Package** | **Language** | **Primary Role** | **Main Inputs** | **Main Outputs** |
|---|---|---|---|---|
| **NBDCtools** | R | Core package for metadata-driven dataset construction and preprocessing | Local tabulated data, metadata, and shadow matrices | Analysis-ready data frames (tibbles) |
| **nbdctools** | Python | Python access layer for dataset construction and metadata-driven assembly | Local tabulated data files and metadata | Polars or pandas DataFrames for downstream analysis |
| **NBDCtoolsData** *(data dependency package)* | R | Data package with the release-specific data dictionaries, levels tables, and sessions tables | N/A | Metadata used by NBDCtools and nbdctools |

### Local file-based input design and metadata assumptions

*NBDCtools* is built around a local, file-based input design. Users first obtain a tabulated NBDC release through an approved data-access platform, such as DEAP or DAP, and store the files in a compliant local or cloud storage environment (Fig. 1B). The data directory is expected to hold one file per table, and the metadata for the corresponding release are provided by *NBDCtoolsData*. Users can then assemble any number of analysis datasets in code from the same release files instead of clicking and downloading a separate extract for each analysis question from the web platforms.

This workflow assumes that NBDC tabulated releases follow a regular structure that can be interpreted through standardized metadata. *NBDCtools* uses the metadata to locate the files that contain the requested variables, determine study-specific identifier columns (join keys) of each table, *participant_id*, *session_id* for ABCD, and *run_id* as well for HBCD. It then retrieves categorical levels and session information, and guides later transformations. The release files are handled as one coordinated collection rather than as unrelated tables. Because the metadata are specific to a release, *NBDCtools* also checks that the data files match the requested release. The check compares the latest session in the data with the latest session expected for that release and stops with an error if they differ. It therefore works as a sanity check for major releases, which usually add a session, but it cannot distinguish minor releases of the same major release, such as ABCD 6.0 and 6.1. Advanced users who need metadata that are not part of *NBDCtoolsData*, for example, for prerelease or corrected data, can register their custom data dictionary, levels table, and sessions table for the current R session. In this design, a dataset is defined entirely in code by the study, the release, and the variable and table names. The same definition can be rerun as an analysis evolves, shared with collaborators, and reported in publications to document how a dataset was derived [13]. This approach complements portal-based workflows, where users rely on point-and-click creation of datasets, one at a time.

### Core R workflow

The core R workflow centers on *create_dataset()*, the main high-level function for building datasets from local NBDC tabulated releases. Instead of requiring users to manually locate, read, and merge source files, *create_dataset()* combines these steps into a single metadata-driven workflow (Fig. 2A). Users specify the study, the release, the path to the local data directory, and the variables or tables of interest, and the function uses release-specific metadata to determine how to assemble the dataset. The convenient wrappers *create_dataset_abcd()* and *create_dataset_hbcd()* preset the study. Datasets can be defined by individual variables, by whole tables, or by a mix of both in the same call. From these arguments, *create_dataset()* identifies the files that hold the requested data, determines the identifier columns, reads the required data, and joins them into a single data frame in memory. Because the whole dataset is held in memory, large requests can exceed the available memory. When assembling ABCD requests of 10,000 or more variables, *create_dataset()* evaluates memory requirements beforehand and halts execution prior to file loading if available memory is below 75% of the estimated memory use, unless users choose to bypass the safeguard.

**Figure 2.**
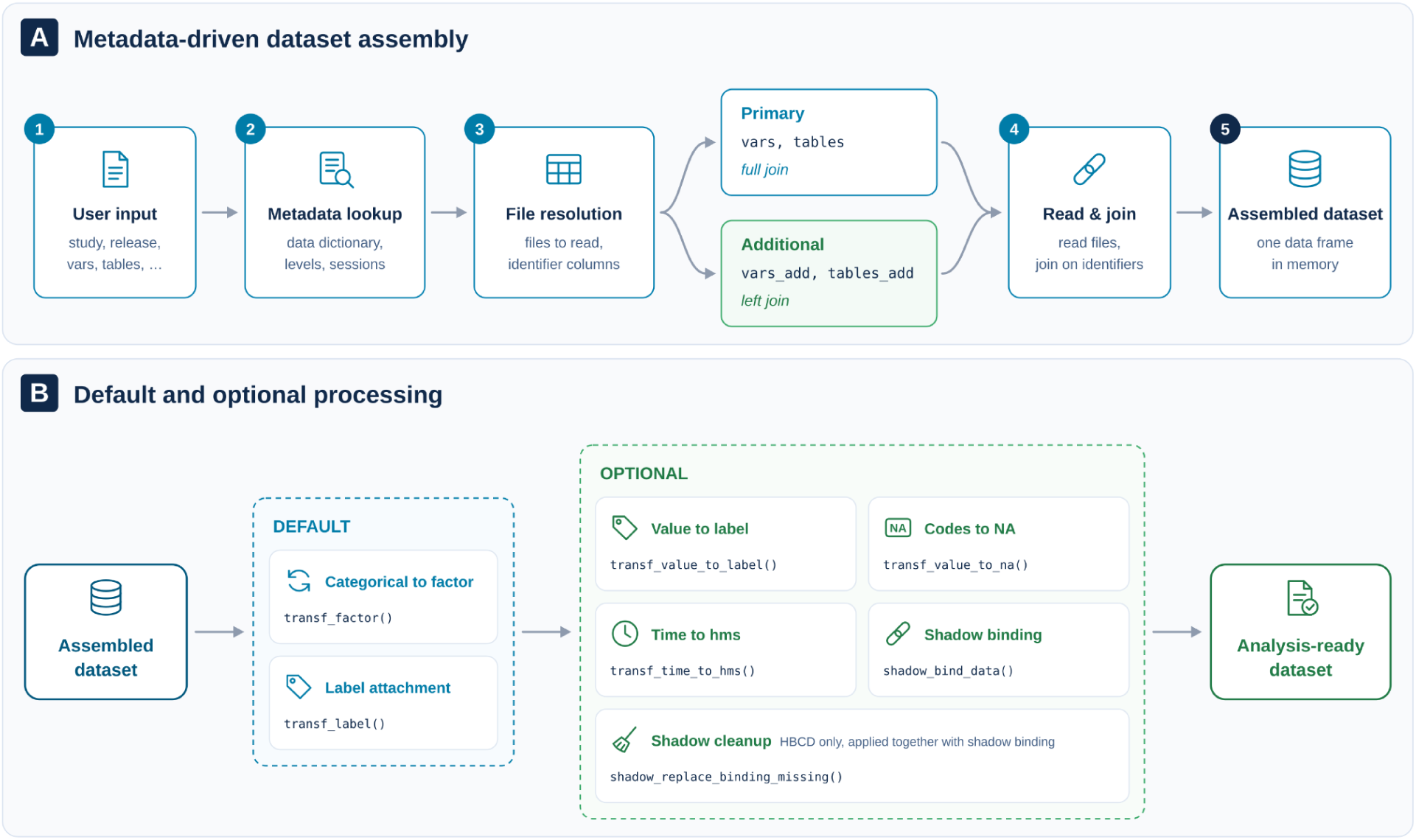
Core dataset-construction workflow in *create_dataset()*. (A) *create_dataset*() uses the user’s input to look up metadata, resolve the files to read, read the data, and join it. Primary selections (*vars*, *tables*) are full joined, and additional selections (*vars_add*, *tables_add*) are left joined to the result. (B) After assembly, default and optional processing steps can be applied within *create_dataset()* or later with the corresponding functions.

After assembly, *create_dataset()* can apply a series of processing steps that make the output more analysis-ready (Fig. 2B). By default, the workflow adds categorical conversion and label attachment steps (Table 2), so variables retain information from the data dictionary and levels metadata. Table 2 lists additional (optional) processing steps that are not turned on by default: (a) convert coded values to their labels (*transf_value_to_label*), (b) map encoded missing-value codes to N/A (*transf_value_to_na*), (c) convert time variables to appropriate time formats (*transf_time_to_hms*), (d) bind shadow-matrix information when available or requested (*shadow_bind_data*), and (e) replace join-induced missingness in shadow matrices (*shadow_replace_binding_missing*). Optional steps can be applied within *create_dataset()* through the corresponding arguments, or by calling these functions directly after dataset assembly. These processing options allow users to maintain a unified data-preparation pipeline while controlling the specific scope and timing of transformation steps according to their analytic requirements.

**Table 2.**
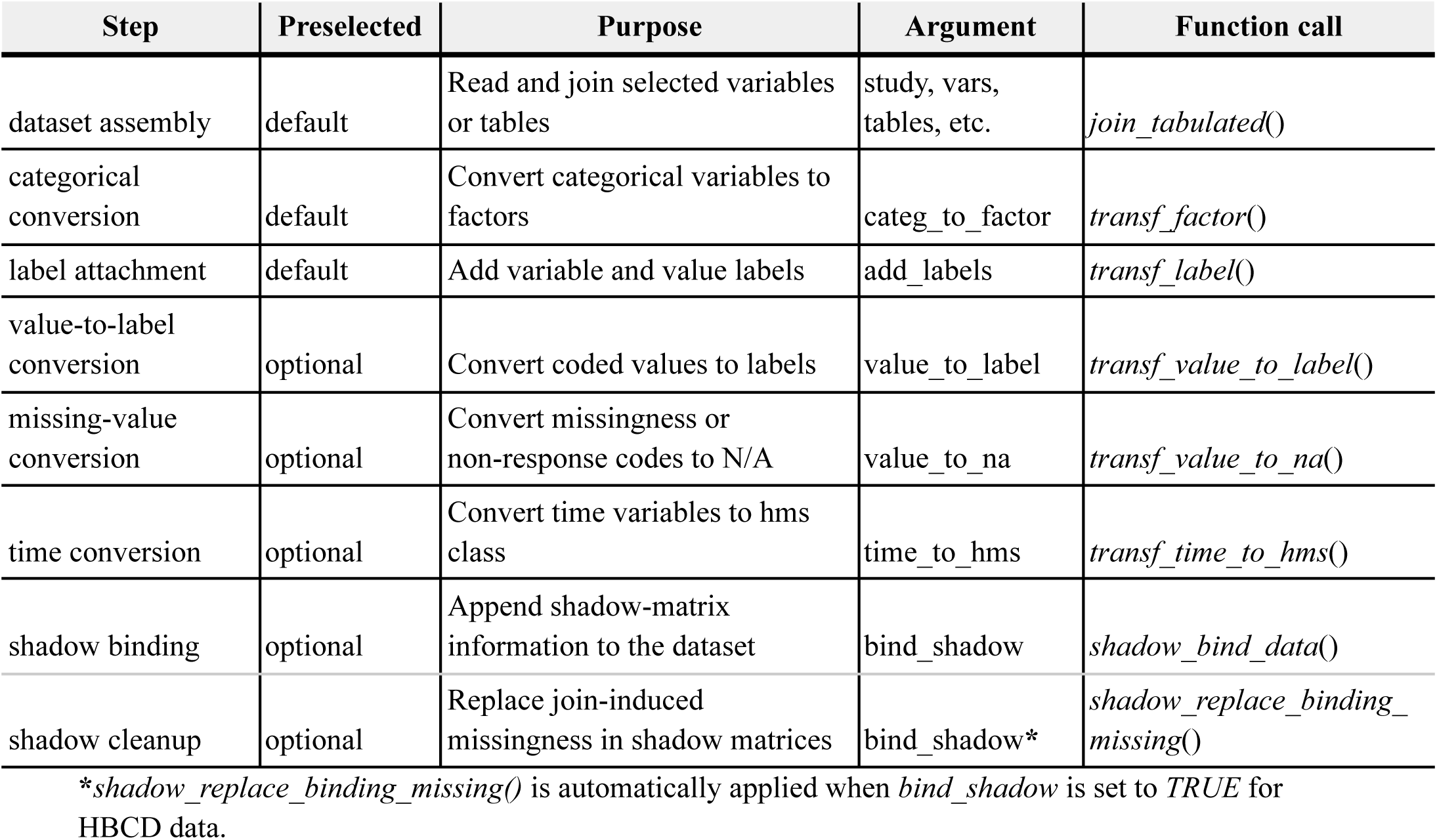
Main processing steps in *create_dataset*(). For each step, the table shows whether it runs by default, the *create_dataset*() argument that controls it, and the function that performs it.

| Step | Preselected | Purpose | Argument | Function call |
|---|---|---|---|---|
| dataset assembly | default | Read and join selected variables or tables | study, vars, tables, etc. | *join_tabulated*() |
| categorical conversion | default | Convert categorical variables to factors | categ_to_factor | *transf_factor*() |
| label attachment | default | Add variable and value labels | add_labels | *transf_label*() |
| value-to-label conversion | optional | Convert coded values to labels | value_to_label | *transf_value_to_label*() |
| missing-value conversion | optional | Convert missingness or non-response codes to N/A | value_to_na | *transf_value_to_na*() |
| time conversion | optional | Convert time variables to hms class | time_to_hms | *transf_time_to_hms*() |
| shadow binding | optional | Append shadow-matrix information to the dataset | bind_shadow | *shadow_bind_data*() |
| shadow cleanup | optional | Replace join-induced missingness in shadow matrices | bind_shadow* | *shadow_replace_binding_missing*() |

**\***\**shadow\_replace\_binding\_missing()* is automatically applied when *bind\_shadow* is set to *TRUE* for HBCD data.

### NBDCtools joining strategy

*NBDCtools* distinguishes between primary and additional selections when joining (Fig. 2A).

Primary variables selected for the main dataset are joined using a full join with identifier columns as the primary key(s), such as *participant_id*, *session_id*, and *run_id*, creating the primary dataset. The additional variables are left joined onto the primary dataset using appropriate identifier columns. In *create_dataset()*, primary variables are specified through *vars* and *tables*, whereas additional variables are specified through *vars_add* and *tables_add*.

This distinction lets users control the shape of the assembled dataset. When variables or tables are part of the primary selection, *NBDCtools* uses a full join to retain all available participant-session records across those primary sources. When variables or tables are added as supplementary information, NBDCtools uses a left join so that these additional fields enrich the existing primary dataset, without adding new rows or join-induced missingness (Fig. S1). This is useful for adding demographic, design, visit-level, or other contextual variables while preserving the analytic structure defined by the main variables of interest. This approach is particularly useful for longitudinal cohort data, where different measures may be collected for different participant-session combinations and where analysts often need to combine focal outcomes with broader contextual variables without introducing unwanted missingness in data. An additional table can be more finely grained than the primary selection, for example when session-level variables are added to participant-level primary variables. Attaching such a table would add rows, so *create_dataset()* stops with an error by default. The argument *finer_add* lets users keep one value per row instead, taken from the earliest available session or run, or keep all values and add the rows, as earlier versions did.

### Processing functions

Beyond the high-level *create_dataset()* workflow, *NBDCtools* provides dedicated, modular processing functions that users can apply to an assembled dataset at any point in an analysis (Table 3). They cover variable transformation, filtering of rows and columns, shadow-matrix handling, metadata access, and utilities for exporting metadata and renaming variables.

**Table 3.** Main function groups in *NBDCtools* beyond *create_dataset()*. The full function reference is available at https://software.nbdc-datahub.org/NBDCtools/reference/index.html.

| Category | Feature | Function |
|---|---|---|
| transformations | convert raw tabulated values into more analysis-ready representations | *transf_factor(), transf_label(), transf_value_to_na(), ...* |
| filtering | subset and clean assembled datasets | *filter_id_events(), filter_events_abcd(), filter_empty_rows(), ...* |
| shadow | bind and clean shadow-matrix information | *shadow_bind_data(), shadow_replace_binding_missing()* |
| metadata | retrieve and customize metadata | *get_metadata(), get_dd(), get_levels(), add_custom_metadata(), ...* |
| utilities | export metadata as BIDS sidecars and convert variable names | *create_bids_sidecar_data(), create_bids_sidecar_metadata(), convert_names_data(), convert_names_file()* |

Transformation functions convert raw tabulated values into analysis-ready representations. Most of these functions can be invoked within *create_dataset()* as default or optional steps (see section Core R Workflow above).

Filtering functions support common cohort-data operations after tables have been joined. Users can filter by participant or event identifiers, apply study-specific event filters, and remove empty rows or columns introduced during multi-table assembly.

The shadow-matrix functions bind a shadow matrix to the data (*shadow_bind_data()*) and, for HBCD, mark cells that are missing only because of a join as “Missing due to joining” (*shadow_replace_binding_missing()*). This separates join-induced missingness from missingness recorded during data collection. Because ABCD does not release shadow matrices, *shadow_bind_data()* can instead derive one from the data with the *naniar* package.

The metadata functions return the data dictionary, levels table, sessions table, identifier columns, and available releases for a study, and *add_custom_metadata()* registers user-supplied metadata.

Two groups of utilities help with data sharing and with changes in variable names across releases. The functions *create_bids_sidecar_data()* and *create_bids_sidecar_metadata()* export the metadata of a dataset or of selected variables as a Brain Imaging Data Structure (BIDS) JSON sidecar file, and *convert_names_data()* and *convert_names_file()* rename variables in a data frame or in a file according to the alternative names recorded in the data dictionary, for example from the names introduced with ABCD release 6.0 to the names used before.

### Python interoperability and pure-Python workflow

The Python package *nbdctools* is intended for users who work mainly in Python. It offers two routes to a dataset. The R-backed interface calls *NBDCtools::create_dataset()* and therefore provides the complete R workflow, whereas the pure-Python implementation reproduces the dataset-assembly logic without R. Python users can run the full workflow from Python, or use the pure-Python path when they need only dataset assembly or cannot install R and its dependencies.

In the R-backed mode (Fig. 3A), users call *nbdctools.create_dataset()* with the same named arguments as the R function, including the study, release, data directory, variables, tables, and processing options. The function passes these arguments through *rpy2* to *NBDCtools::create_dataset()* and returns the result as a *Polars* or *pandas* DataFrame. Four return modes determine how the R data frame is transferred to Python. Two of them convert it directly through *rpy2* (polars, pd). The other two, the sink modes, write it to a temporary Arrow IPC (Feather) file with the R package *arrow* and read that file into Python (polars_sink, pd_sink). Python users therefore get the same joining and preprocessing behavior as R users and can continue their analysis in Python.

**Figure 3.**
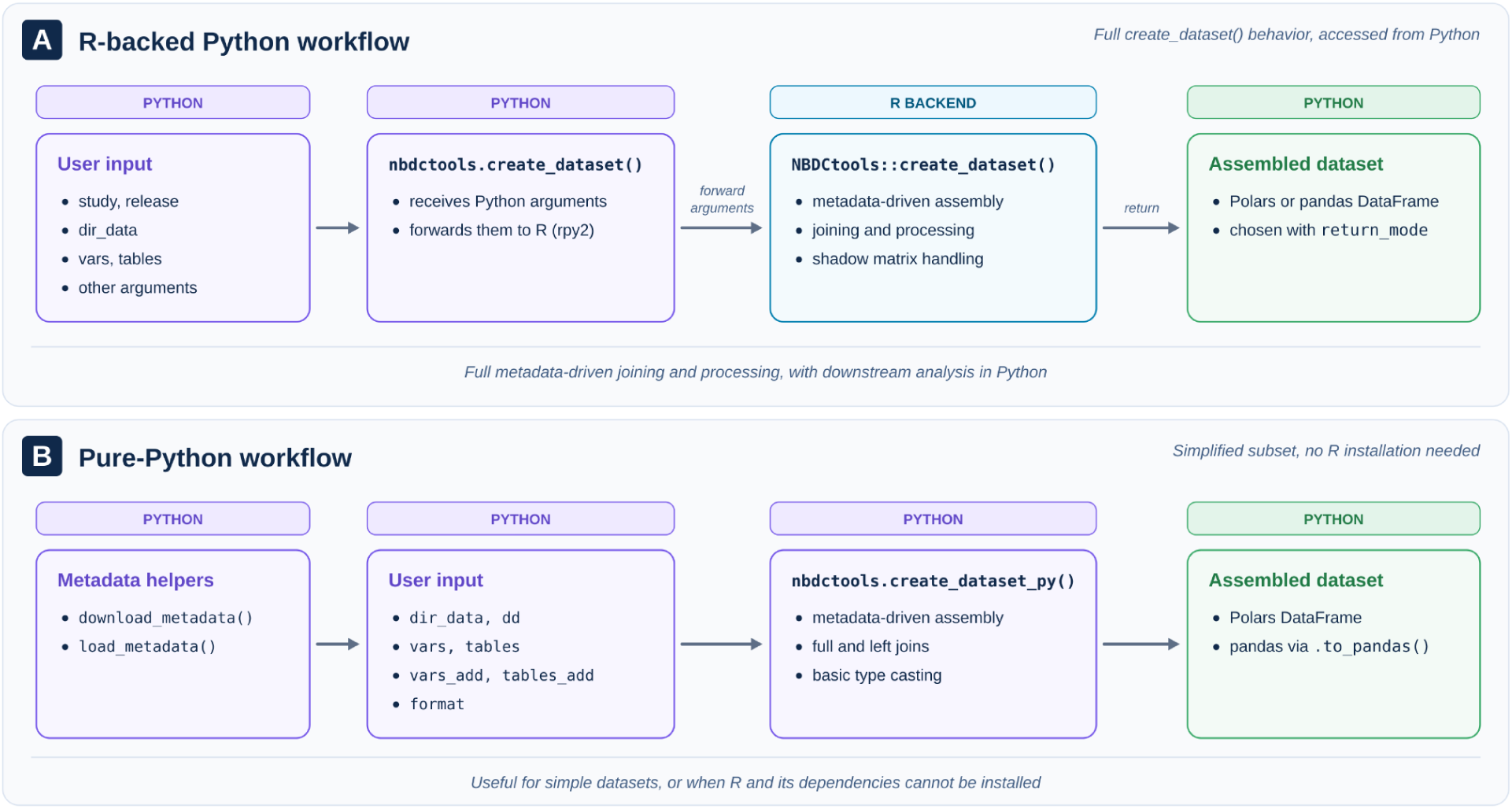
Python workflows in *nbdctools*. (A) In the R-backed workflow, *nbdctools.create_dataset()* forwards named arguments to *NBDCtools::create_dataset()* and returns the assembled dataset as a *Polars* or *pandas DataFrame*. (B) In the pure-Python workflow, the metadata helpers *download_metadata()* and *load_metadata()* and the function *create_dataset_py()* assemble a dataset without R. The result is a *Polars DataFrame*, which can be converted to *pandas*.

The pure-Python workflow, *nbdctools.create_dataset_py()*, assembles datasets directly from the release files and the metadata without R (Fig. 3B). The metadata are downloaded once with *download_metadata()* and loaded with *load_metadata()*. The function selects the files based on the data dictionary, combines primary selections with full joins and additional selections with left joins, casts columns to the data types recorded in the data dictionary, and converts nominal and ordinal variables to categorical columns. Additional selections never add rows. If an additional table is more finely grained than the primary selection, the function keeps one value per row and issues a warning. It returns a *Polars* DataFrame. Labels, conversion of missingness codes, time conversion, removal of empty rows, shadow matrices, and the release check are not supported, so this path is a simplified alternative to the R implementation rather than a replacement for it. Users who need those steps can use the R-backed interface.

## Results

### Overview

The following sections present *NBDC Data Tools* from three angles. We first apply the workflow to a cross-domain ABCD example in R and Python (Fig. 4). We then show how the dataset definition from that example can be reused when the analysis, the collaborators, the release, or the programming language change (Fig. 5). Finally, we report benchmarks of runtime and memory use for dataset construction in R and for the Python interfaces (Figs. S2 and S3).

**Figure 4.**
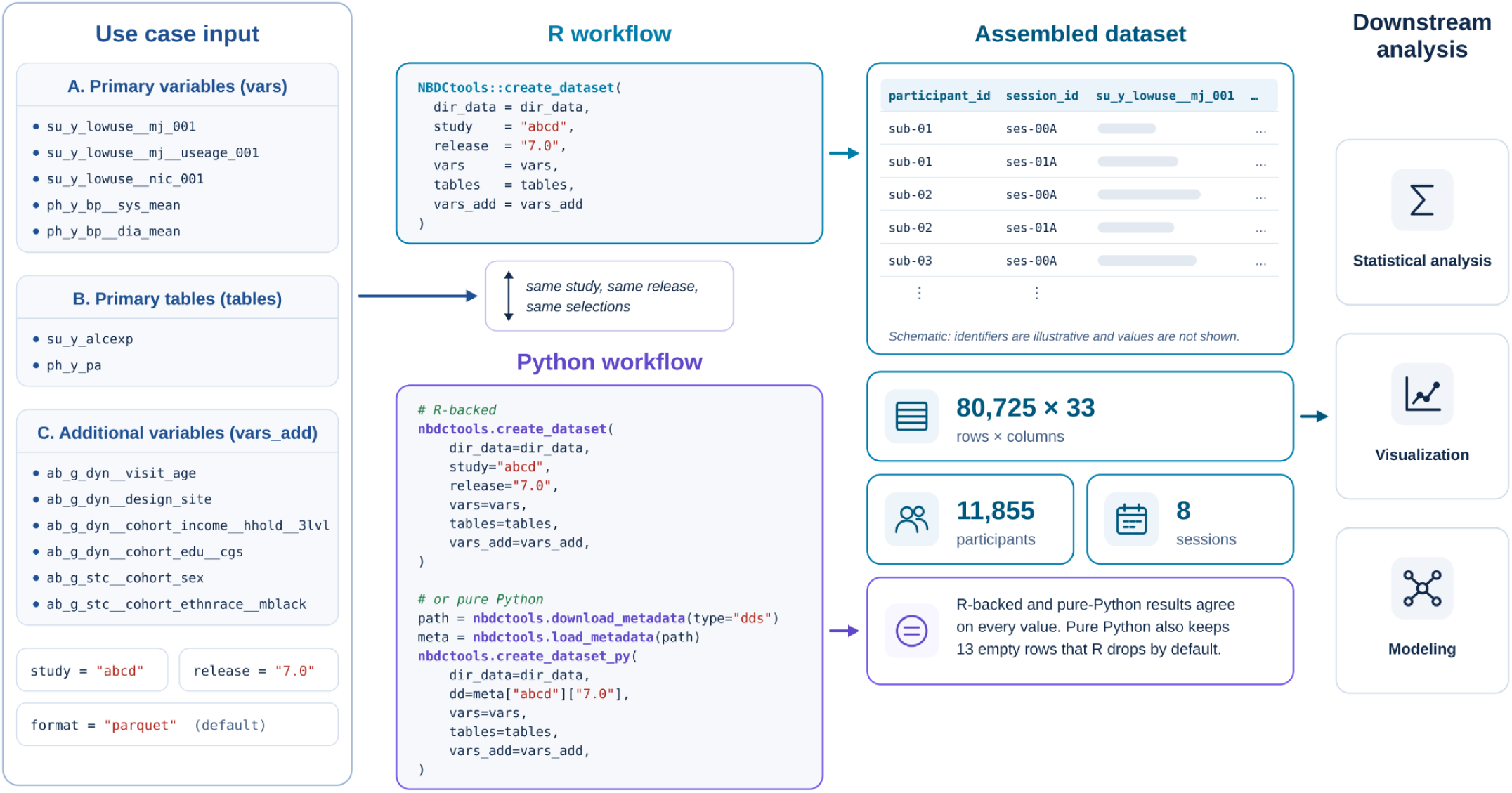
Cross-domain use case. A dataset on youth substance use and physical health is assembled from the ABCD 7.0 tabulated release with the same arguments in R and in Python. The R call and the R-backed Python call returned 80,725 rows × 33 columns. The pure-Python call matched these rows value for value and returned 13 extra rows in which all primary variables were missing. The R function removes such rows by default. The assembled dataset is the starting point for statistical analysis, visualization, and modeling.

**Figure 5.**
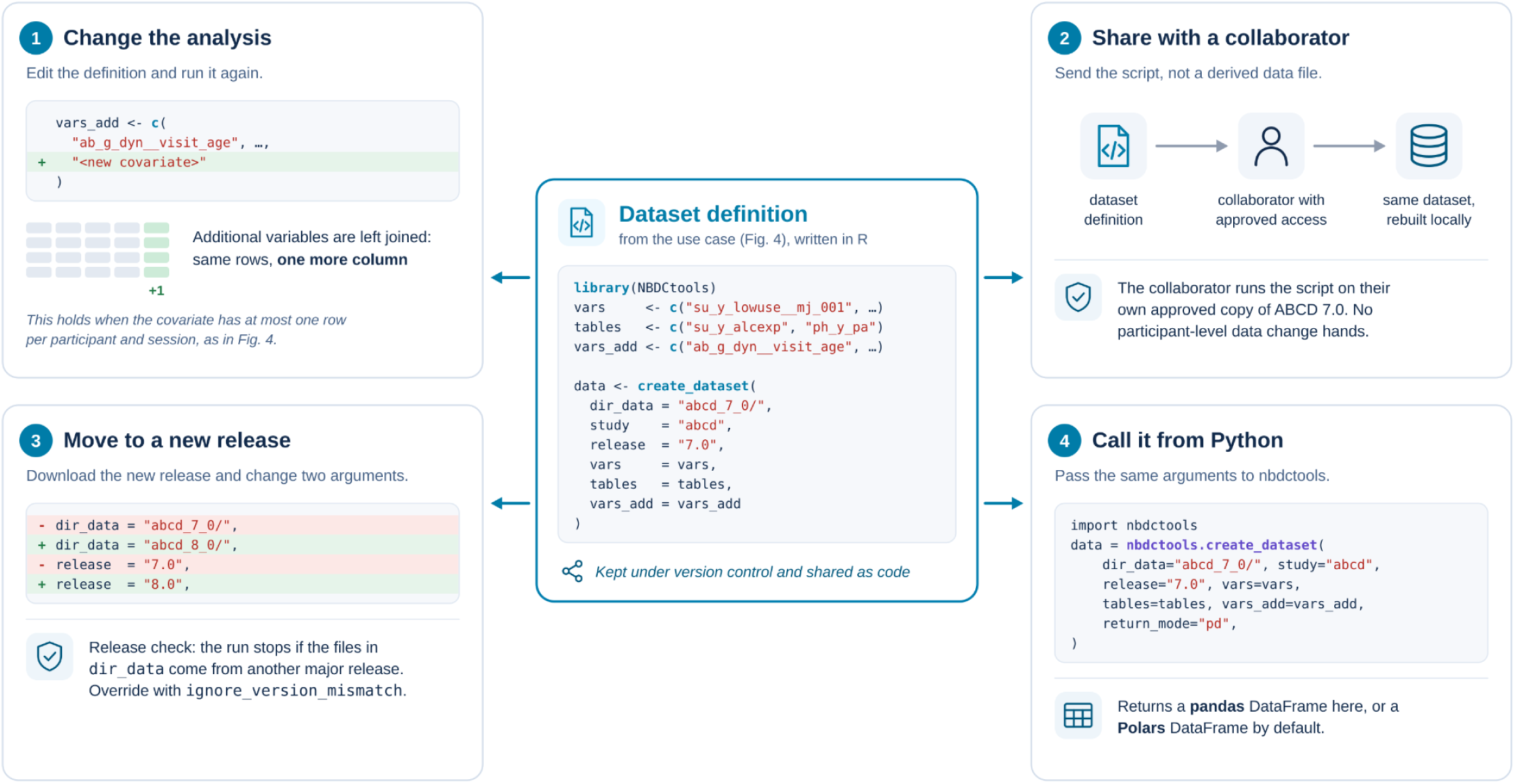
Reusing one dataset definition. The definition from the use case (center) is rerun after a covariate is added, shared with a collaborator who runs it on their own approved copy of the release, moved to a new release by changing the *release* argument, and called from Python with the same arguments. In each case, only code is exchanged, and the dataset is rebuilt from the original release files.

### Use case

To show the workflow in a realistic setting, we built a cross-domain dataset from the ABCD 7.0 tabulated release in R and in Python. The dataset was designed for studying associations between youth substance use and physical health. We chose this example because it combines several common tasks in one call, namely selecting individual variables, selecting whole tables, attaching covariates as additional variables, and joining data across domains (Fig. 4).

The definition (Fig. 4) selected five primary variables (*vars*), three on youth marijuana and nicotine use from the low-level substance-use table (*su_y_lowuse__mj_001*, *su_y_lowuse__mj__useage_001*, *su_y_lowuse__nic_001*) and two blood-pressure summaries from the youth blood-pressure table (*ph_y_bp__sys_mean*, *ph_y_bp__dia_mean*). It also selected two whole tables (*tables*), one on youth alcohol expectancies (*su_y_alcexp*) and one on youth physical activity (*ph_y_pa*). Six demographic and design covariates were added through *vars_add*, including visit age, assessment site, household income, caregiver education, sex, and ethno-racial identity (*ab_g_dyn__visit_age, ab_g_dyn__design_site, ab_g_dyn__cohort_income__hhold__3lvl, ab_g_dyn__cohort_edu__cgs, ab_g_stc__cohort_sex, ab_g_stc__cohort_ethnrace__mblack*). The last two come from the static table *ab_g_stc*, so they were joined on *participant_id* alone and repeated across each participant’s sessions.

In R, *NBDCtools::create_dataset()* built a dataset of 80,725 rows and 33 columns from the local ABCD 7.0 release, covering 11,855 participants across 8 sessions. Passing the same arguments to *nbdctools.create_dataset()* returned the same number of rows and columns. Given the same selections, the pure-Python *create_dataset_py()* matched that result value for value in all 80,725 rows. The only difference was 13 extra rows in which all primary variables were missing, which the R function removes by default (Fig. 4). The number of rows is large relative to the number of participants because the full join of the primary sources keeps every session at which any of the selected measures was collected. An analysis restricted to, for example, the annual visits would first filter these rows with *filter_events_abcd()*.

### Reusing a dataset definition

In *NBDC Data Tools*, the dataset definition, not a data file, is what users keep under version control and share. Figure 5 shows four situations in which the definition from the use case is reused. When the analysis changes, users edit the definition, for example by adding a covariate to *vars_add*, and run it again. Because additional variables are left joined, the rebuilt dataset gains a column and keeps its rows. When a collaborator wants to reproduce the dataset, the authors share the script rather than a derived data file, and the collaborator runs it on their own approved copy of the same release. When a new release becomes available, users download it, point the data directory to the new files, and change the *release* argument. The release check described in the Methods stops the script if the files come from a different major release. Finally, the same arguments can be passed from Python to *nbdctools.create_dataset()*, which returns the dataset as a *Polars* or *pandas* DataFrame. In all four cases, only code changes hands, and the dataset is rebuilt from the original release files.

### Benchmark

Because NBDC releases contain hundreds of tables and tens of thousands of variables, we measured how runtime and memory use scale with the size of a request. Benchmark code and additional results are available in the package documentation (https://software.nbdc-datahub.org/NBDCtools/articles/benchmark.html and https://software.nbdc-datahub.org/nbdctools-py/guide/benchmark/).

The R benchmark used the ABCD 6.0 release on a Linux server (Intel Xeon Gold 6448H, 4 TB RAM) that read the data from a network drive at about 350 MB/s. We timed *join_tabulated()*, the function that reads and joins the data inside *create_dataset()*, for requests built from randomly selected whole tables (30 to 700 tables) or from randomly selected variables, up to about 80,000 variables, with three replications per request (Fig. S2). Runtime depended mainly on how many tables had to be read to supply the requested variables. At about 40,000 variables, selecting whole tables took roughly 40 minutes, whereas drawing the same number of variables at random from across the release took roughly 2 hours (Fig. S2A). Memory use was similar for both strategies and grew linearly with the number of variables, from about 15.8 GB for 10,000 variables to about 125 GB for 80,000 variables (Fig. S2B). Converting categorical variables to factors with *transf_factor()* added about 11%, for example, 17.6 GB instead of 15.8 GB at 10,000 variables (Fig. S2C). Peak memory during processing is higher than the size of the final data frame because data are copied during joining and transformation, so we recommend having two to three times that size available. For most users, these figures are conservative. Local solid-state drives read data much faster than the network drive used here, and typical analyses select far fewer variables. The 33-column dataset of the use case, for example, fits easily in the memory of a laptop.

The Python benchmark compared the four return modes of *nbdctools.create_dataset()* with the pure-Python *create_dataset_py()* and with a baseline in which *NBDCtools::create_dataset()* ran in a fresh R process and returned only the dimensions of the result, so the baseline transferred no data to Python (Fig. S3). We ran the benchmark on ABCD 7.0 on a laptop (Intel Core i7-13700H, 32 GB RAM, solid-state drive) with a request for 8 tables and 12 variables, which produced 139,095 rows and 825 columns with every method. Each method was timed three times after one untimed warm-up run, and every run started in a fresh process. All R-backed methods received the same arguments, with labels turned off and the release and memory checks bypassed. Direct conversion through *rpy2* was slow, with median runtimes of 172.5 s for conversion to *Polars* and 28.6 s for conversion to *pandas*, compared with 15.6 s for the baseline. The sink modes took 16.0 s (*Polars*) and 15.5 s (*pandas*), within 0.4 s of the baseline. The pure-Python path finished in 3.7 s, about 4.2 times faster than the baseline. It does less work, however, because it only joins the data and casts data types (see Methods).

Two practical recommendations follow from these results. Requests run faster when they draw on fewer tables, so selecting whole tables, or limiting a request to the tables an analysis needs, reduces runtime. In Python, the sink modes should be preferred for all but very small datasets, provided that enough temporary disk space is available for the intermediate file.

## Discussion

### Main features and contribution

*NBDC Data Tools* gives researchers a local, scripted way to turn the per-table files of an NBDC tabulated release into datasets ready for analysis. The release-specific metadata coordinates variable selection, table assembly, identifier handling, and downstream processing. The design of *NBDC Data Tools* addresses a prevalent informatics challenge inherent in large-scale longitudinal research by enabling the systematic assembly of analysis-ready datasets from high-dimensional releases in a way that others can inspect and repeat. Because the dataset definition is code, the script that produces a dataset also documents how it was derived, and the same definition runs in R and in Python. This matters most when datasets have to be rebuilt often, carried over to new releases, shared with collaborators, and described in publications.

### NBDC Data Tools and web-based data platforms

*NBDC Data Tools* complements the web platforms of the NBDC Data Hub and does not replace them. The web platforms handle authorization and are where users discover, explore, and download NBDC data. NBDC Data Tools starts after these steps, once an approved user has downloaded a tabulated release and wants to build datasets in a local R or Python environment (Table 4).

**Table 4.** Roles of NBDC Data Tools and web-based data platforms in preparing analysis datasets. ✓ indicates direct support and ◐ partial or indirect support. A blank cell means that the feature is not a primary role of the tool.

| **Feature** | **NBDC Data Tools** | **Web platforms** |
|---|---|---|
| Scriptable dataset construction in R/Python | ✓ |  |
| Reusable dataset definition in code | ✓ |  |
| Metadata exploration | ✓ | ✓ |
| Primary and additional variable selection | ✓ | ✓ |
| Mix full-table and variable selection | ✓ | ✓ |
| Cross-release workflow reruns | ✓ | ◐ |
| Produces analysis-ready outputs for downstream analysis | ✓ | ◐ |
| Data visualization and custom plotting |  | ◐ |
| Interactive data discovery and browsing |  | ✓ |
| Approved data access and download |  | ✓ |

The two approaches suit different stages of a project. Web platforms work well for browsing the data dictionary, exploring variables, and creating a dataset interactively. A programmatic definition pays off once a dataset has to be rebuilt. Changing the variable list or moving to a new release then means editing a line of code and running it again, rather than repeating a series of selections and downloading a new file. Only one copy of each release needs to be stored, because every analysis dataset is rebuilt in memory from the same files instead of being saved as a separate extract. Most importantly, the definition is declarative and runs on the original release files. A collaborator with approved access can therefore rebuild the same dataset from the script, whereas an analysis script shared on its own often fails because it starts from a derived data file that only its author has.

### Limitations

NBDC Data Tools has several limitations. It is not a data-access system. Users must already have approved access and must have downloaded the tabulated release to a local or other approved computing environment, and the web platforms remain necessary for authorization, discovery, and download. The workflow also depends on the metadata. *NBDCtoolsData* has to be updated for each new release, and errors in a released data dictionary or levels table carry over into the assembled datasets. The software assumes that releases keep the regular structure described by the metadata, with one file per table and consistent identifier columns, and it currently supports only the ABCD and HBCD studies. Because datasets are assembled and held in memory, very large requests need a machine with correspondingly large memory. Finally, the packages need continued maintenance as the studies, the release formats, and the underlying R and Python libraries change.

### Future work

We are developing *NBDCvalidation*, a companion R package for quality control of NBDC tabulated data (https://github.com/nbdc-datahub/NBDCvalidation). It checks the files, metadata, data values, and shadow matrices of a dataset against NBDC standards and summarizes the results in a report. Our release teams that curate NBDC releases run these checks on every data delivery. Errors in files, metadata, or values then surface before the data reach researchers, and every release is held to one set of criteria. External users can apply the checks to their own data in NBDC format, such as datasets they pair with custom metadata through *add_custom_metadata()*, before analyzing or sharing them.

The suite will also follow the development of the NBDC Data Hub. *NBDCtoolsData* will be updated with each new ABCD and HBCD release, and other NBDC studies that follow the same metadata conventions could be supported by adding their metadata. The Data Hub is also preparing cloud-based analysis environments, and we plan to make NBDC Data Tools available there so that the same scripts can be used.

## Conclusion

*NBDC Data Tools* lets researchers build datasets from NBDC tabulated releases with a few lines of R or Python code. The release-specific metadata determines which files are read, how tables are joined, and how values are converted, and the code that defines a dataset documents how it was derived. As the NBDC Data Hub adds releases and studies, the same scripts can be rerun on new data, which makes the step from approved data access to statistical analysis easier to repeat and to reproduce.

## Software and data availability

The packages of *NBDC Data Tools* are open source and released under the GNU General Public License (version 3 or later). They can be installed from public package repositories and from GitHub, where the source code is available. Documentation and tutorials are available on the package websites. At the time of writing, the current versions are *NBDCtools* 1.1.2, *NBDCtoolsData* 6.0.0, and *nbdctools* 1.0.3. ABCD and HBCD data are available to approved researchers through the NBDC Data Hub (https://www.nbdc-datahub.org/). No participant data are shared with this paper.

NBDCtools

- Installation: CRAN, GitHub (https://github.com/nbdc-datahub/NBDCtools).
- Website: https://software.nbdc-datahub.org/NBDCtools/

NBDCtoolsData

- Installation: r-universe, GitHub (https://github.com/nbdc-datahub/NBDCtoolsData).
- Website: https://software.nbdc-datahub.org/NBDCtoolsData/, or see NBDCtools

nbdctools

- Installation: PyPI, GitHub (https://github.com/nbdc-datahub/nbdctools-py).
- Website: https://software.nbdc-datahub.org/nbdctools-py/

## Acknowledgement

We would like to acknowledge the contributions of the ABCD Study® consortium and HBCD consortium, particularly the experts who offered valuable input and guidance throughout the development of the *NBDC Data Tools*. Additionally, we extend our gratitude to the participants of both studies for their time and commitment to advancing science.

## References

1. Feldstein Ewing S, Luciana M. The adolescent brain cognitive development (ABCD) consortium: rationale, aims, and assessment strategy [special issue]. Dev Cogn Neurosci. 2018;32: 1–164.

2. Walhovd KB, Fjell AM, Westerhausen R, Nyberg L, Ebmeier KP, Lindenberger U, et al. Healthy minds 0-100 years: Optimising the use of European brain imaging cohorts (“Lifebrain”). Eur Psychiatry. 2018;50: 47–56. doi:10.1016/j.eurpsy.2017.12.006

3. Nelson CA, Frankeberger J, Chambers CD. An introduction to the HEALthy Brain and Child Development Study (HBCD) study. Dev Cogn Neurosci. 2024;69: 101441. doi:10.1016/j.dcn.2024.101441

4. Linkersdörfer J, Berman S, Smith C, Ziemer L, Zhang L, Abad S, et al. Strategies for collection, management, and release of data for multi-site longitudinal studies: Lessons from the ABCD Data Analysis, Informatics, & Resource Center. Dev Cogn Neurosci. 2026;80: 101754. doi:10.1016/j.dcn.2026.101754

5. Jernigan TL, Brown SA, Dowling GJ. The adolescent brain cognitive development study. J Res Adolesc. 2018;28: 154–156. doi:10.1111/jora.12374

6. Saragosa-Harris NM, Chaku N, MacSweeney N, Guazzelli Williamson V, Scheuplein M, Feola B, et al. A practical guide for researchers and reviewers using the ABCD Study and other large longitudinal datasets. Dev Cogn Neurosci. 2022;55: 101115. doi:10.1016/j.dcn.2022.101115

7. Li L, Bayat M, Hayes TB, Thompson WK, Neale MC, Gard AM, et al. Missing data approaches for longitudinal neuroimaging research: Examples from the Adolescent Brain and Cognitive Development (ABCD) Study®. Dev Cogn Neurosci. 2025;74: 101563. doi:10.1016/j.dcn.2025.101563

8. Zhang L, Celhay O, Das B, Berman S, Ziemer LR, Smith CJ, et al. ABCDscores: an R package for computing summary scores in the ABCD Study®. Front Neuroinform. 2026;20: 1858950. doi:10.3389/fninf.2026.1858950

9. Wilson G, Bryan J, Cranston K, Kitzes J, Nederbragt L, Teal TK. Good enough practices in scientific computing. PLoS Comput Biol. 2017;13: e1005510. doi:10.1371/journal.pcbi.1005510

10. Lopez DA, Cardenas-Iniguez C, Subramaniam P, Adise S, Bottenhorn KL, Badilla P, et al. Transparency and reproducibility in the Adolescent Brain Cognitive Development (ABCD) study. Dev Cogn Neurosci. 2024;68: 101408. doi:10.1016/j.dcn.2024.101408

11. Hanscombe KB, Coleman JRI, Traylor M, Lewis CM. ukbtools: An R package to manage and query UK Biobank data. PLoS One. 2019;14: e0214311. doi:10.1371/journal.pone.0214311

12. McCarthy P. funpack. Zenodo; 2026. doi:10.5281/zenodo.20701883

13. Wilkinson MD, Dumontier M, Aalbersberg IJJ, Appleton G, Axton M, Baak A, et al. The FAIR Guiding Principles for scientific data management and stewardship. Sci Data. 2016;3: 160018. doi:10.1038/sdata.2016.18

