## Supplement fig 1-3 for "NBDC Data Tools: Building Analysis-ready Datasets From ABCD and HBCD Releases in R and Python"

### Supplement for NBDC Data Tools

#### Figures

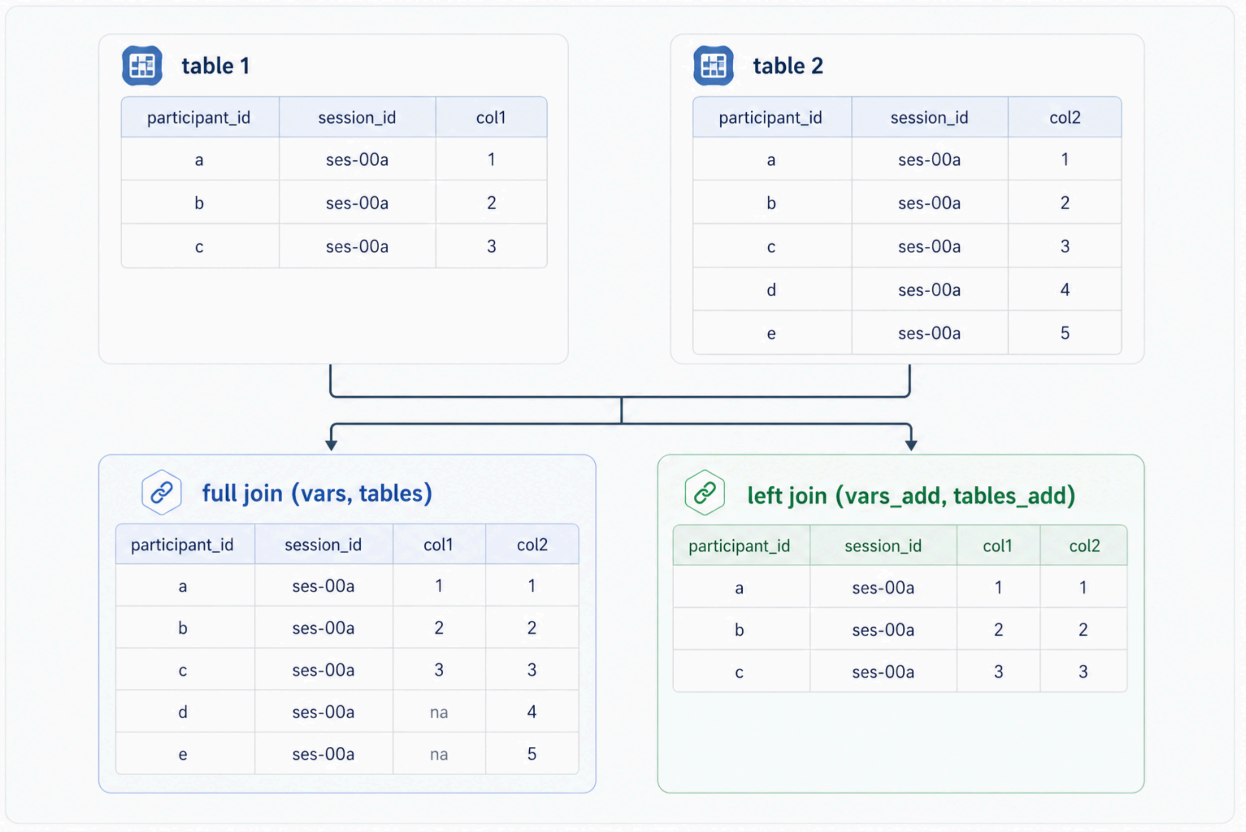


**Figure S1**. Schematic illustration of full and left joins in *NBDCtools*. The example shows how *NBDCtools* handles primary and additional selections during dataset assembly. For the left join case, *table 1* is used as the main table. Primary selections specified through *vars* and *tables* are full joined, and additional selections are left joined.

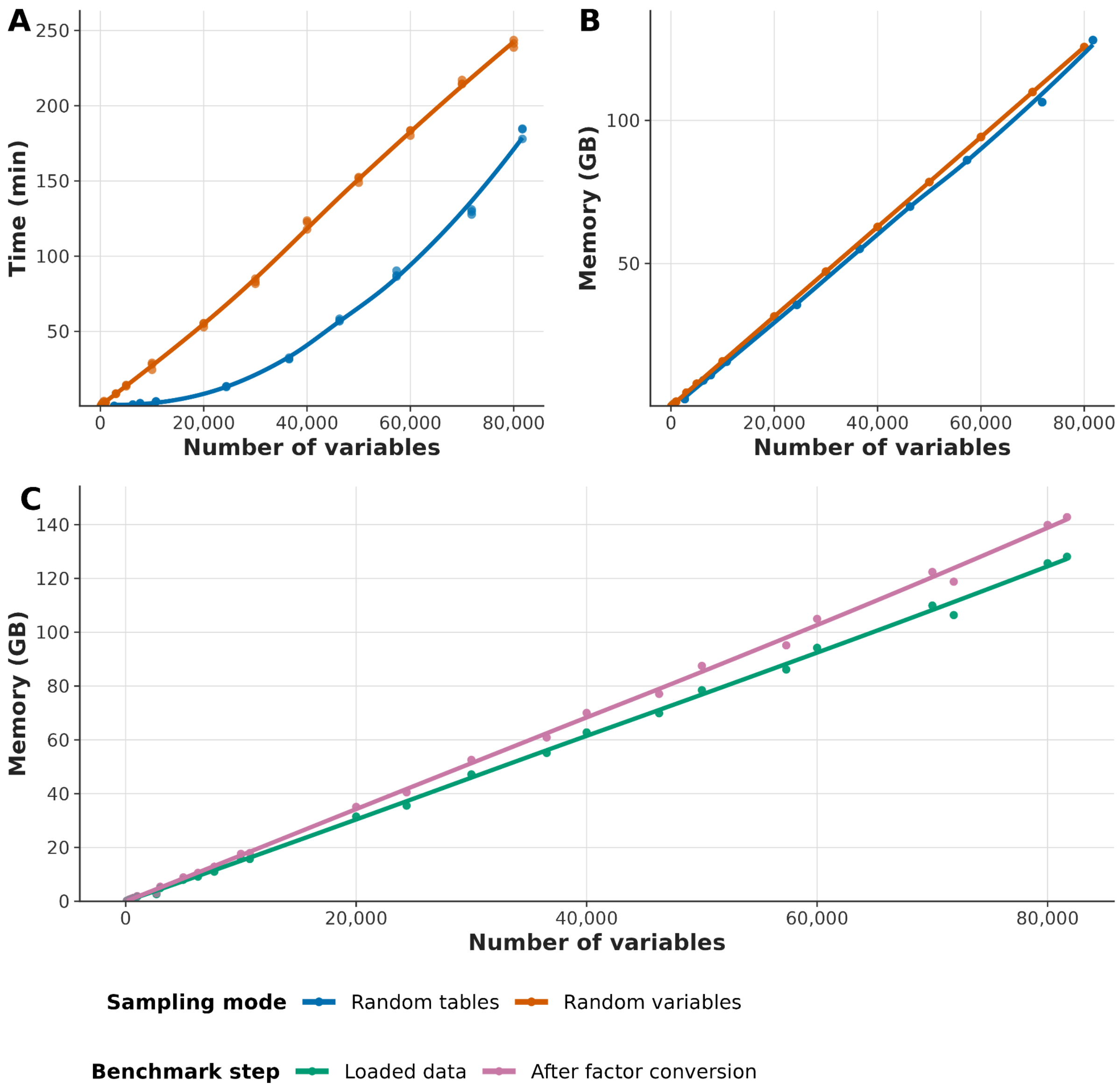


**Figure S2**. Benchmark performance of *NBDCtools* data loading and processing. (A) Time required to load and join tabulated data with join_tabulated() as a function of the number of variables, comparing randomly selected tables with randomly selected variables. (B) Memory usage for the same loading step. (C) Memory usage before and after the *additional* transf_factor() processing step. Points show individual benchmark runs, and lines show LOESS-smoothed trends. Memory is reported in GB and time in minutes.

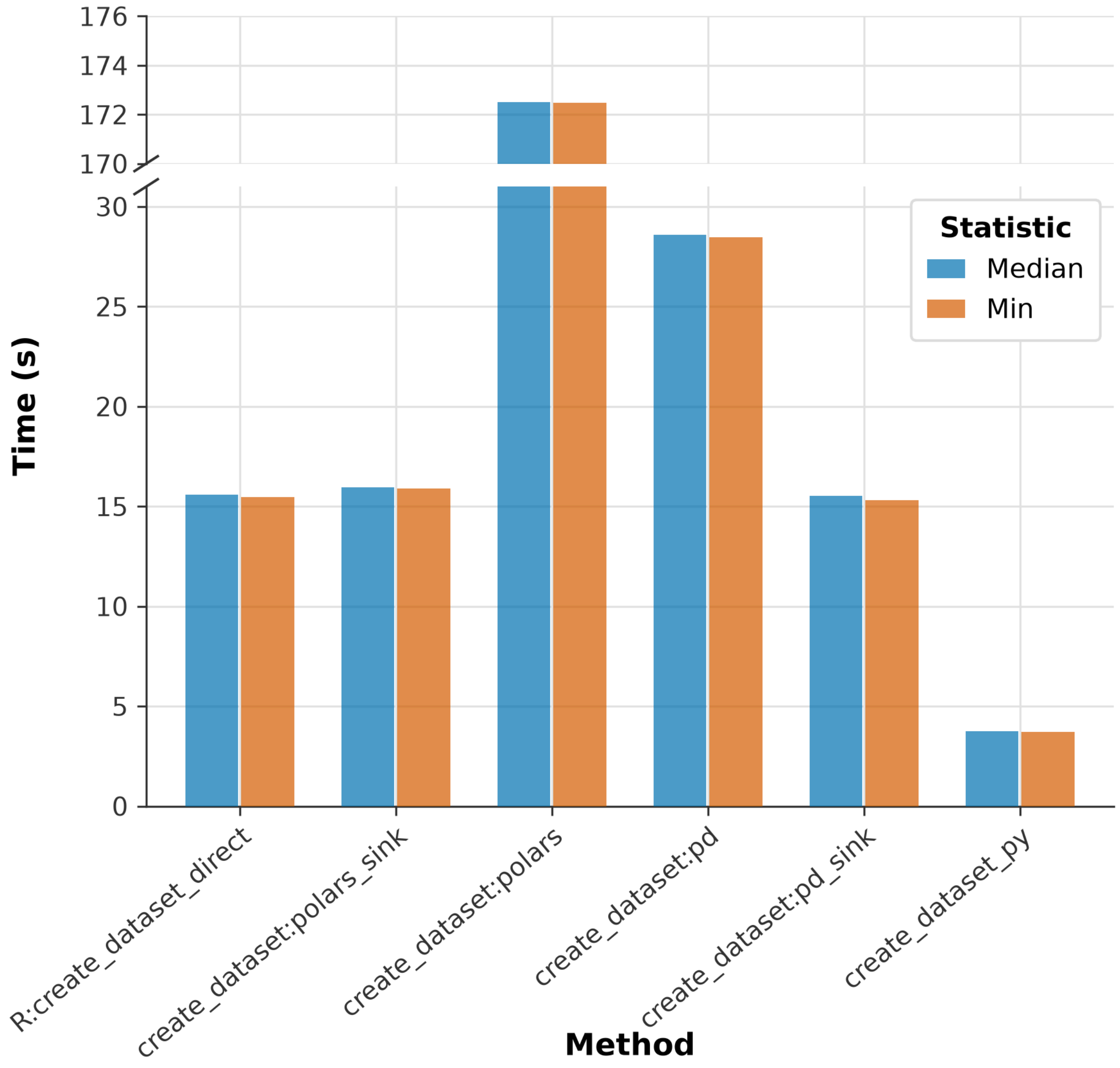


**Figure S3**. Benchmark of dataset construction methods comparing minimum and median runtimes (seconds) in *nbdctools* on the ABCD 7.0 release (139,095 rows × 825 columns). Bars show the median and minimum runtime of each method over three runs, each in a fresh process. The y-axis is broken between 31 s and 170 s so that the slowest method (172.5 s) can be shown with the others (3.7 to 28.6 s). *R:create_dataset_direct* calls *NBDCtools::create_dataset* in a fresh Rscript process and returns only the dimensions of the result, so this baseline transfers no data to Python. *create_dataset:polars_sink* and *create_dataset:pd_sink* use the wrapper, write the result to a temporary Arrow IPC (Feather) file, and read it back into Polars or pandas. *create_dataset:polars* and *create_dataset:pd* convert the R data frame directly through rpy2. *create_dataset_py* is the pure-Python implementation.
